# Predicting Individual Traits from Unperformed Tasks

**DOI:** 10.1101/2021.10.12.464045

**Authors:** Shachar Gal, Niv Tik, Michal Bernstein-Eliav, Ido Tavor

## Abstract

Relating individual differences in cognitive traits to brain functional organization is a long-lasting challenge for the neuroscience community. Individual intelligence scores were previously predicted from whole-brain connectivity patterns, extracted from functional magnetic resonance imaging (fMRI) data acquired at rest. Recently, it was shown that task-induced brain activation maps outperform these resting-state connectivity patterns in predicting individual intelligence, suggesting that a cognitively demanding environment improves prediction of cognitive abilities. Here, we use data from the Human Connectome Project to predict task-induced brain activation maps from resting-state fMRI, and proceed to use these predicted activity maps to further predict individual differences in a variety of traits. While models based on original task activation maps remain the most accurate, models based on predicted maps significantly outperformed those based on the resting-state connectome. Thus, we provide a promising approach for the evaluation of measures of human behavior from brain activation maps, that could be used without having participants actually perform the tasks.

## Introduction

Cognitive abilities and psychological traits widely differ between individuals. The search for objective measures of individual traits is a long-lasting challenge for the scientific community, spanning psychological questionnaires, behavioral tests and more recently, brain imaging-derived measures (Haynes and Rees, 2006). Specifically, it has recently been shown that individual scores of general intelligence could be predicted using individuals’ task-induced brain activation patterns, acquired while performing a task inside a magnetic resonance imaging (MRI) scanner (i.e., task-functional MRI, hereafter referred to as task-fMRI) (Greene et al., 2018; Sripada et al., 2020). However, tasks used in fMRI studies are often tedious and time-consuming, require participants cooperation and should be pre-designed to target the cognitive domain of interest. What if, alternatively, individual traits could be predicted from ‘task’ activation maps without the need to actually perform any task?

Resting-state (rs) fMRI is a procedure in which no explicit task is introduced to participants while they are scanned (Biswal et al., 1995).The rs-fMRI signal can be analyzed to extract patterns of functional connectivity, reflecting time-synchronous activity of spatially distinct brain areas commonly represented as the rs-connectome (Sporns, 2011). In the last few years, the rs-connectome was found to be stable enough to be used as a fingerprint to detect identity (Finn et al., 2015), to correlate with behavioral and demographic measurements (Smith et al., 2015) as well as personality traits (Cai et al., 2020; Dubois et al., 2018a), and was also linked to individuals’ genetic profile (Colclough et al., 2017). Functional connectivity fingerprinting and the prediction of individual measures from resting-state connectivity are methods under constant development (Kashyap et al., 2019; Kong et al., 2019; Shen et al., 2017). However, rs-connectome-based predictions of intelligence were less successful than predictions based on task-fMRI data (Gao et al., 2019; Greene et al., 2018; Sripada et al., 2020). A widespread explanation for this is that placing the brain in a more cognitively demanding state (e.g., task vs. rest) improves brain-based prediction of intelligence in a way analogous to treadmill testing of cardiac function (Greene et al., 2018; Sripada et al., 2020).

Still, brain activity and connectivity are tightly linked. Patterns of functional connectivity show a striking spatial correspondence with various task-induced brain activation patterns (Smith et al., 2009), and features of functional connectivity were shown to accurately predict task-induced activations on an individual basis across a wide range of cognitive domains (Cole et al., 2016; Tavor et al., 2016). In the present work we aim to predict individual-specific cognitive measures, such as general intelligence, using task-free functional MRI scans. Our hypothesis is that task activation maps predicted from resting-state connectivity patterns (hereafter referred to as “connTask” maps), can be used to predict individual traits. Furthermore, we hypothesize that predictions of individual traits using models based on these “connTask” maps would be more accurate than predictions derived from models based on the rs-connectome directly. Thus, we suggest a novel approach for accurate estimation of cognitive abilities from a simple and effortless fMRI scan, without actually performing any task.

To test our hypotheses, we used the data of 847 participants from the Human Connectome Project (HCP) who underwent resting-state and several task-fMRI scans, as well as multiple behavioral tests performed outside of the scanner (see methods) (Barch et al., 2013; Glasser et al., 2013; Smith et al., 2013; Van Essen et al., 2012). As a measure of intelligence, we calculated individual general cognitive ability (G) scores based on 10 cognitive tests, using a previously suggested factor analysis approach (Dubois et al., 2018b). We used a 10-fold cross-validation approach to generate a variety of connTask maps, and predict individual G-scores using a Brain Basis Set modeling paradigm (Sripada et al., 2020, 2019a) (see methods and Figure 1). We predicted intelligence scores using either original task activation maps, connTask maps or resting-state functional connectomes, and compared the success of predictions derived from these different inputs. In addition, we devised models that combine data from multiple task-activation/connTask maps and compared their success to models based on a single map (see methods).

**Figure 1.**
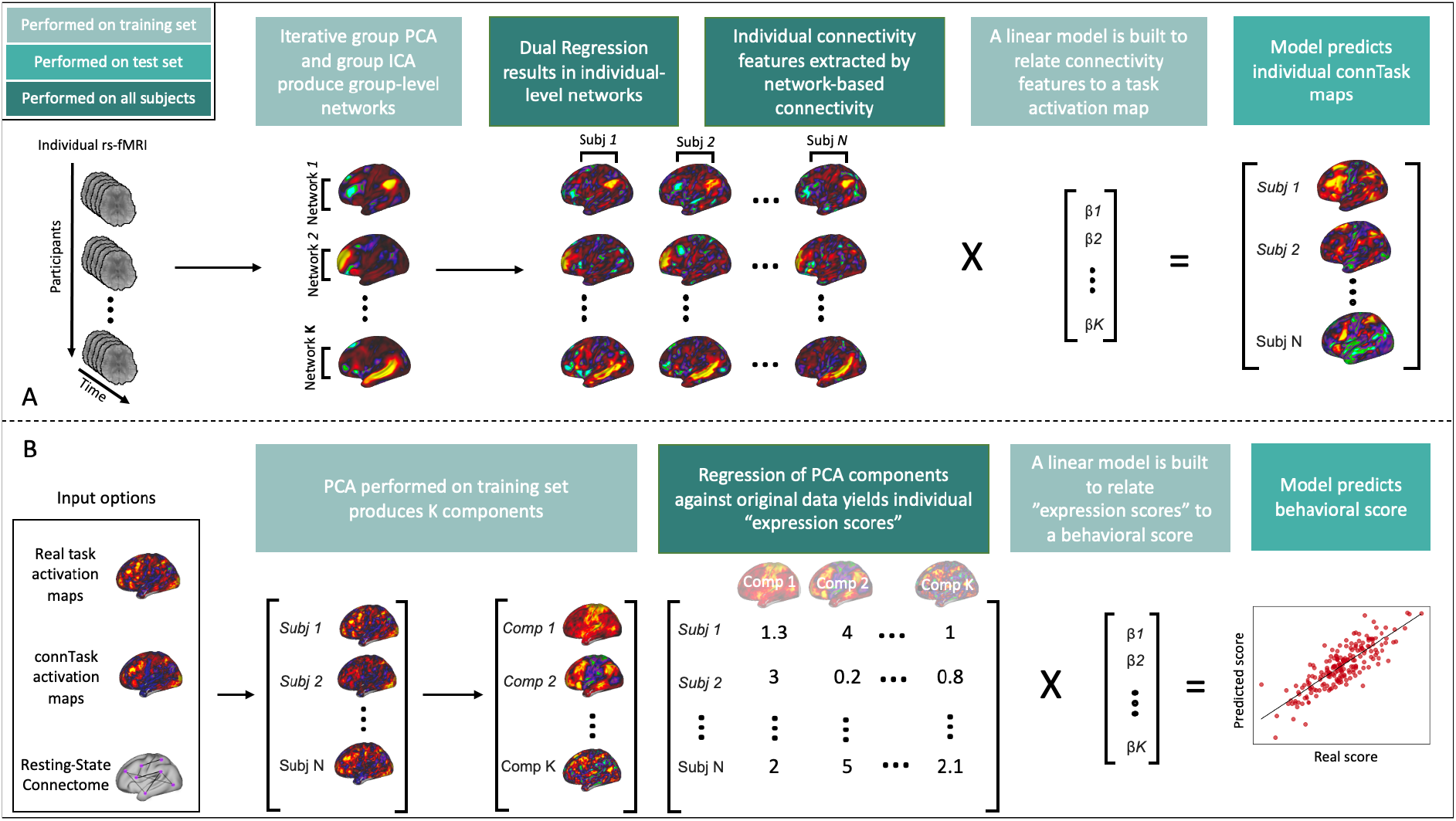
Data processing pipelines. **A)** connTask prediction pipeline from rs-fMRI time series to individual task-induced brain-activation (adapted from Tavor et al., 2016) **B)** Individual trait prediction pipeline using Brain Basis Set modeling (Sripada et al., 2020, 2019a). Colors encode the subset of the sample on which the procedure was performed.

## Materials and Methods

### Data and participants

We used the data from the Human Connectome Project (HCP) (Van Essen et al., 2012) for all subsequent analyses. The dataset includes functional and structural minimally pre-processed (Glasser et al., 2013) scans (1200 subjects release).

We included participants that performed all the relevant behavioral and cognitive tests (see G-score construction, section 2.6), and completed all the resting-state and task fMRI sessions, barring the Motor task, which was left out of further analysis due to its weak contribution to the prediction of intelligence (Sripada et al., 2020), and the low specificity of its connTask maps (Tavor et al., 2016). These criteria resulted in a dataset of 847 participants.

All acquisition parameters and image processing pipelines are described in detail in Glasser et al (2013). Briefly, all fMRI data (rest and task alike) were scanned with a TR of 0.72s. Each participant included in this study had four resting-state fMRI runs with a total of 1,200 timepoints per run, and performed all necessary tasks fMRI scans.

As part of the HCP’s pre-processing procedure, all functional data were de-noised using FMRIB’s ICA-based Xnoiseifier (FIX) (Griffanti et al., 2014; Salimi-Khorshidi et al., 2014), which identifies independent components of structured artefacts in the data, and regresses them out. This is the standard method for nuisance regression in the HCP. This procedure ensures that the “cleaned” data is minimally affected by noise sources such as white-matter signal or motion effects, as these are represented as noise components and are regressed out of the data. The “cleaned” data was then resampled and “projected” onto a surface representation consisting of 91,282 “grayordinates” in standard space. Data were aligned and registered using Multimodal Surface Matching (MSMAll) (Robinson et al., 2018).

As for the task fMRI, data provided by the HCP were already post-processed and included statistical analysis. All tasks in the HCP were performed in a block-design manner, and activity estimates were computed for the time series from preprocessed functional scans using FSL’s FILM (FMRIB’s Improved Linear Model with autocorrelation correction) (Woolrich et al., 2001). “Blocks” of each type of stimuli included in each task, were convolved with a double gamma canonical hemodynamic response function (Glover, 1999) to generate the main model regressors.

All tasks were performed twice, and the statistical maps provided by the HCP are the result of the second level analysis, that provides a mean result for each contrast across both runs.

The Working Memory task included 405 time-points, with 25 seconds long blocks, and 4 blocks per condition (0-back, 2-back); The Gambling task included 253 time-points, with 28 seconds long blocks, and 2 blocks per condition (reward, punish); The Language task included 316 timepoints, with blocks of varying length (average ~30 seconds, math and story blocks were maintained at same length), and 4 blocks per condition (math, story); The Social task included 274 time-points, with 23 seconds long blocks and 2 or 3 blocks per condition (run 1 contained 2 Social and 3 Random blocks and Run 2 contained 3 Social and 2 Random blocks.); The Relational task included 232 time-points with 16 seconds long blocks, and 3 blocks per condition (relational, control); The Emotion task included 176 time-points, with 18 seconds long block and 3 blocks per condition (face, shape).

We used the Z-score maps provided by the HCP for all contrasts described in Table 1 with no further analysis. Additional details regarding task design and processing can be found in (Barch et al., 2013)

**Table 1.** Task contrasts used in further analyses. Contrasts in bold were used as exemplars in the main analysis. Additional contrasts were used in the analyses presented in supplementary figures S8 and S12.

| Task Domain | Contrasts |
| --- | --- |
| Working Memory | 2bk>0bk |
|  | **2bk** |
|  | 0bk |
| Language | **Math-Story** |
| Social | **Random** |
|  | TOM |
|  | TOM-Random |
| Relational | **Rel** |
|  | Match |
| Gambling | **Punish** |
|  | Reward |
| Emotion | **Faces-Shapes** |

### Construction of connectivity-derived task activation (connTask) maps

Connectivity-derived task activation maps were calculated according to the method suggested by Tavor and colleagues (Tavor et al., 2016), for which all codes are publicly available. The procedure is described in Figure 1A and detailed below.

#### Cross-validation

ConnTask maps were predicted using the following procedure in a10-fold cross validation routine, where in each iteration 9/10 of the participants were used as the training set and the remaining participants were used as the test set. Considering the fact that the HCP dataset includes siblings, we made sure that in each iteration of the cross-validation process, participants who are genetically related were allocated to the same group (train/test), as to prevent over-fitting (Colclough et al., 2017).

#### Feature extraction

The feature extraction procedure included four steps, designed to yield a set number of functional connectivity maps, to be used in the prediction model.

1. First, in order to reduce the dimensionality and prepare the data for the following steps, all the training set’s pre-processed resting-state fMRI data were combined using an iterative group principal component analysis (PCA) procedure (Smith et al., 2014). This step is aimed to concatenate a large number of participants; an iterative PCA was shown to yield a very accurate approximation to a concatenation of the whole dataset while having very low memory requirements (Smith et al., 2014). In each iteration of this procedure, data from one restingstate scan (1200 timepoints) were added, and the overall dimensionality was reduced to a set of 1000 group-level components.
2. Group independent component analysis (ICA) (Beckmann et al., 2005) was performed on the training set’s reduced data using FastICA (Hyvärinen, 1999), yielding 45 spatially-independent cortical components to be used as seeds for connectivity analysis. Five of these components were manually classified as noise according to their spatial maps and removed from further analyses (see Supplementary Figure S1).
3. Dual regression was performed on the cortical components against individual time-series, to produce subject-specific cortical independent component (IC) maps (Beckmann et al., 2009) for all participants, test and training sets alike. In dual regression, the first regression uses the cortical group-ICA maps as regressors to get an individual time-series for each component per participant. The second regression uses the individual time-series as regressors to get individual spatial maps. Thus, this step produces individual seeds to be used in the following connectivity analysis.
4. Last, the subject-specific IC maps were used as seeds in a weighted seed-to-vertex analysis. As such, individual IC maps were regressed against individual rs-fMRI time-series in order to yield one time-course per spatial map. Each time course was then correlated with the original rs-fMRI data to produce an individual connectivity map for each IC map. In the original work by Tavor and colleagues, weighted seed-to-vertex connectivity maps were created also for additional 32 sub-cortical components. Of these, we included 3 cerebellar components that were created through group-ICA. The remaining sub-cortical components had little to no contribution to the prediction success, and thus were kept out of the procedure we implemented in this work.

#### Model fitting and prediction

A general linear model was used to map the functional connectivity features to task data (i.e. individual z-score contrast maps derived from fMRI task analysis). For each task activation map (i.e., a specific contrast of a specific task), a specific model was fitted. As performed in Tavor et al., 2016, all models were broken down spatially into 50 non-overlapping regions of interest. Regions were defined according to the resting-state data, using ICA, and allocating vertices to parcels using a majority-vote on the ICA result. Within each of these 50 parcels, a general linear model was used to relate connectivity features to activation data, using the training set’s data. Prior to model fitting, all features were normalized and standardized across the whole brain.

### Construction of rs-connectomes

For each participant, the time-series from all 4 runs of rs-fMRI scans were normalized, concatenated and demeaned. We used a parcellation that separates the cortex into 360 nonoverlapping areas (Glasser et al., 2016), and calculated the averaged resting-state time-series within each parcel. Pearson’s correlation coefficients were computed between each pair of parcels.

### Further processing of the data

All fMRI data were scaled and normalized before being used for prediction, using scikit-learn’s (Buitinck et al., 2013) StandardScaler. Original task-activation maps were masked to include only cortical data, in order to make them comparable with the connTask data, where activity is predicted only for the cortex. Removing the subcortical areas from the original task-activation data has actually improved the prediction results compared with whole-brain maps.

### Trait prediction procedures

#### Cross-validation

Similar to the prediction of the connTask maps, the prediction of individual traits was also performed using a 10-fold cross validation routine, taking into account the family structure in the HCP data. In each iteration, a model was trained on 9/10 of the data, and predictions were yielded for 1/10 of the data, while genetically related participants are kept in either the training or the test datasets. Importantly, data splits were identical in the two cross-validation procedures (e.g., connTask maps prediction and trait prediction) meaning that for each 1/10 of the data, connTask maps were predicted using the remaining 9/10 of the data, and also trait prediction was based on the same 9/10 of the data.

#### Brain Basis Set prediction pipeline

We utilized the Brain Basis Set (BBS) prediction pipeline proposed by Sripada and colleagues (Sripada et al., 2020, 2019a) using multiple functionalities provided by the Scikit-Learn python package (Buitinck et al., 2013).

The BBS prediction pipeline is composed of 4 steps:

A. The training sets’ fMRI data is reduced using Principal Components Analysis (PCA) to a predetermined number of components ***K***.
B. These components are regressed against the individual data of every participant, to yield a number we refer to as an “expression score”.
C. The training sets’ expression scores are used to fit a linear model that predicts the desired trait.
D. The model is applied on the test set.

These steps are visualized in figure 1B.

This pipeline was used for all predictions performed on a single input (i.e., original task activation maps, connTask maps or resting-state connectomes).

#### Pipeline adaptation for multiple inputs

We further adapted the pipeline to enable a combination of features extracted from multiple inputs (i.e., task/connTask maps). The leading notion in devising this procedure was to prevent an inflation of the number of features used in the regression model as we add more map inputs.

With this notion in mind, we set a total number of components, to be extracted from all inputs combined (***T***, for total), which is kept uniform across all types of combinations (e.g., the same ***T*** is used when combining 2 maps or 6). Once ***T*** is set, steps A+B are performed for each input separately, with ***K*** being defined as 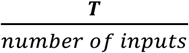. Thus, a sum of ***T*** “expression scores” is calculated for each subject.

We then reduce the number of features from ***T*** to a predetermined number, ***F***, by correlating each “expression score”, across participants, to the predicted trait, and choosing the top ***F*** correlated scores. Steps C+D are then performed using the remaining ***F*** features, with the minor change of using a regularized linear model, to better handle the larger number of features.

We fitted an Elastic Net regression model with an L1 ratio of 0.001, which is almost equivalent to ridge (L2) regression. The penalty factor was determined in a nested 5-fold cross validation process (i.e., a separate cross validation performed in each iteration on the training set folds only). We used Scikit-Learn’s ElasticNetCV.

#### Pipeline parameters

We implemented the single-input BBS pipeline with a ***K*** of 75, as in Sripada et al (Sripada et al., 2020, 2019b; Taxali et al., 2021).

We implemented the expanded, multi-input, BBS pipeline with a ***T*** (i.e., total number of components extracted across inputs) of 300 and an ***F*** (i.e., number of features after reduction using correlation analysis) of 160.

The value for ***T*** was chosen as to ensure that even when a model is based on a combination of up to 6 different inputs (as in the largest combination tested in this study), the number of components extracted from each single input will not be less than 50. The Value for ***F*** was chosen by examining 20 different values, ranging between 50 to 250 (see Supplementary Figure S2).

### Construction of the G-score

We conducted an exploratory factor analysis, following the procedure suggested by Dubois and colleagues (Dubois et al., 2018) and the code made available by them.

Participants were included in this analysis if they completed all relevant tasks, and their MMSE (Mini-Mental State Examination) score was greater than 26. This left a total of 1,181 participants. Unadjusted scores from ten cognitive tasks were used in the analysis, including seven tasks from the NIH Toolbox (Dimensional Change Cart Sort, Flanker Task, List Sort Test, Picture Sequence Test, Picture Vocabulary Test, Pattern Completion Test, Oral Reading Recognition Test), and three tasks from the Penn Neurocognitive Battery (Penn Progressive Matrices, Penn Word Memory Test, Variable Short Penn Line Orientation Test). This means that some of the participants included in this analysis are not included in any further analyses, due to lacking MRI data (whether rest or task). However, their inclusion here increases the statistical power of modeling the latent intelligence factor (g).

We performed the exploratory factor analysis in a 10-fold cross validation procedure, such that the g-scores for each 1/10 of the participants were produced using weights calculated on the data of the other 9/10 of the participants.

### Assessment of significant differences in prediction results

In order to assess whether there is a significant difference between the accuracy of the predictions based on two different data-sets (e.g., connTask map and rs-connectome), we performed 1000 iterations where we split the data randomly to training and test sets (9/10 train, 1/10 test, family structure taken into consideration), and predicted behavioral scores. In every iteration, prediction success for each data-set was measured by Pearson’s r between actual and predicted scores. Significant differences in prediction success were detected using a non-parametric, related samples test (Wilcoxon signed ranks test), which takes into account the overlap in training sets across iterations (Demšar, 2006).

We compared prediction success of models based on 14 different contrasts, or combinations of contrasts, to the prediction success of a model based on resting state connectivity. We performed this comparison using original task maps as well as connTask maps, which resulted in a total of 28 comparisons. Therefore, all p values were corrected using Bonferroni correction for multiple comparisons, for 28 comparisons.

The comparison of predictions of in-scanner and out-of-scanner measurements followed a similar procedure. While the main analysis compared values of prediction success from two different models predicting the same target score, in this analysis we compared the ratio or amount by which the models based on original task activation maps outperformed models based on connTask maps, in various traits.

### Assessment of significance of prediction success against a null distribution

We assessed model prediction success significance using a permutation test. To create the “null distribution” against which we then compare our result, we ran each model 10,000 times with the target variable (i.e., the cognitive score we wish to predict) being shuffled for each iteration. We recorded the prediction success from each of those iterations, which represents the “null distribution” of prediction success scores, and assigned the real model a p value of 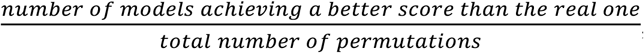, meaning that the minimal possible p value is 0.0001.

This p value was compared to a Bonferroni corrected to 29 comparisons, as we added a comparison for the resting-state connectome.

### Construction of model contribution maps

In order to visualize the distribution of predictive information in each task contrast, we multiplied each component derived from the data reduction step of the BBS pipeline (see step A in the BBS prediction pipeline above) by its corresponding beta coefficient from the linear model (see step D). These weighted components were then summed, within and across the 10 folds, to yield the model contribution maps. The process is described by Equation (1):

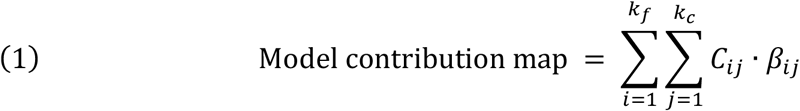

Where *k_f_* is the number of fold used in the cross-validation routine, *k_c_* is the number of components yielded in the dimensionality reduction step (step A in the BBS pipeline), *J* is the *j^th^* components in the *i^th^* iteration, and *β_ij_* is the beta coefficient assigned to the “expression score” associated with *J* in the prediction model (step D in the BBS pipeline).

### Quantification of meaningful contribution from each rs-network

In order to quantify the contribution of each of the seven canonical resting-state networks (Yeo et al., 2011) to the prediction model, we used the HCP’s workbench command (Marcus et al., 2011) function “wb_command -cifti-find-clusters” to find clusters of meaningful contribution in the model contribution maps. We set the surface-value-threshold to the 0.99 percentile of each map’s values (“contribution power” in arbitrary units) and the surface-minimum-area to 15 mm^2^. We then counted the number of significantly contributing clusters in each of the resting-state networks, separately for the prediction from original and connTask maps and for the prediction from single vs. multiple contrasts.

## Results

### Prediction of intelligence scores

As hypothesized, connTask-based models performed significantly better than the model based on rs-connectome (V < 0.0001, Bonferroni corrected for 28 comparisons; Figures 2–3, and Supplementary Table S2). All models yielded predictions that were significantly more accurate than chance, as asserted by a permutation test (V < 0.005, Bonferroni corrected for 28 comparisons). For both original and connTask data, models that combined features extracted from multiple maps resulted in an improved prediction. Group-level prediction accuracy was generally higher using original task activation maps rather than predicted ones. However, for each task there was a considerable subset of participants (ranging from 39% to 49%) for which predictions based on the connTask maps were even more accurate than those based on original task maps (see Supplementary Figures S3-4).

**Figure 2.**
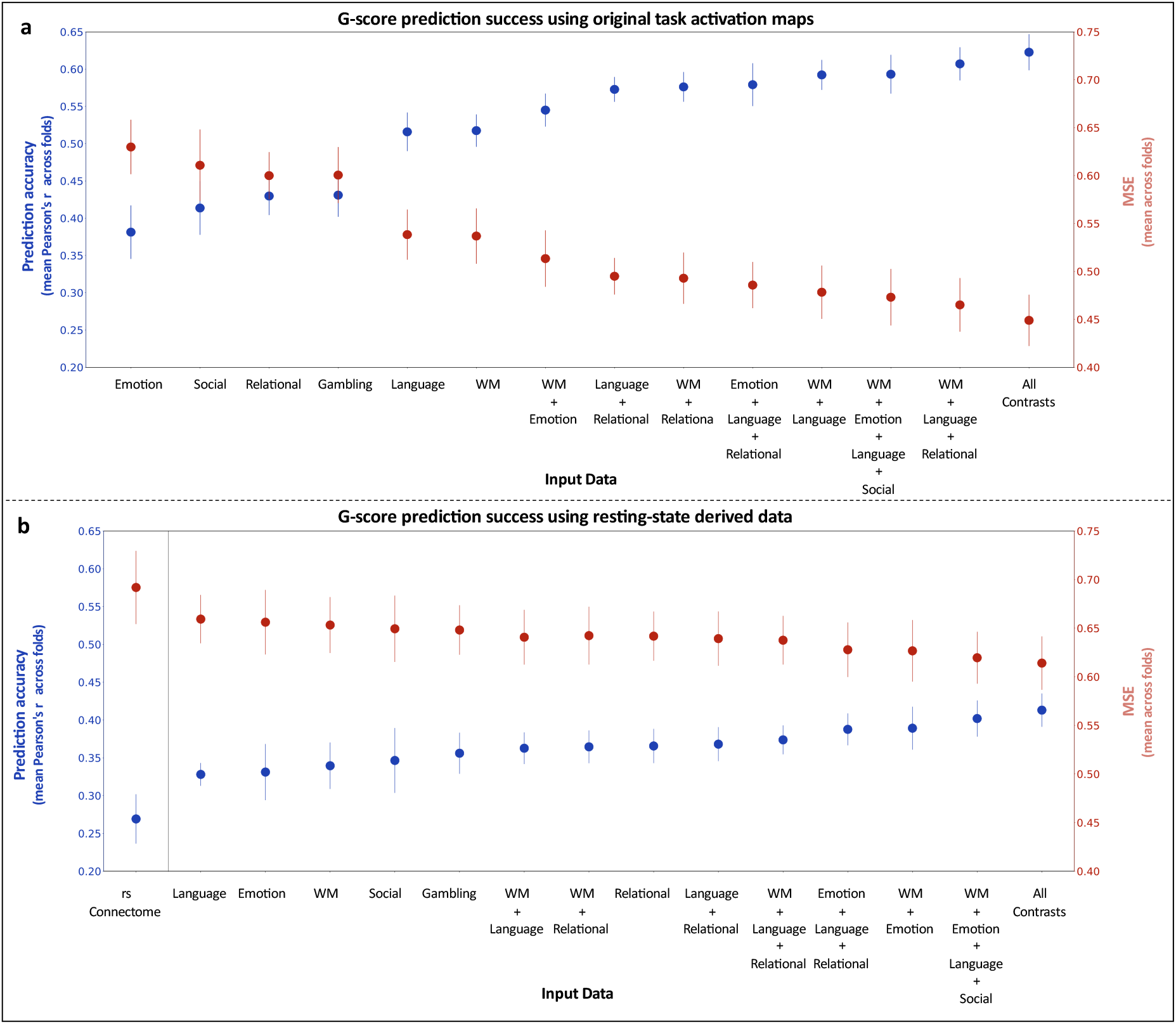
G-score prediction success of models based on real activation maps, predicted activation maps and the rs-connectome. Prediction success was estimated by the Pearson correlation between predicted and observed G-scores (in blue, referring to the left y-axis) and the mean squared error of the prediction (in red, referring to the right y-axis). Error bars depict standard error of the measurement between cross-validation folds. **A)** Prediction success of models based on original task activation maps. **B)** Prediction success of models based on resting-state fMRI derived data (rs-connectomes and connTask maps). All connTask-based models were significantly more accurate than the rs-connectome based model (*P* < 0.0001, Bonferroni corrected for 28 comparisons).

**Figure 3.**
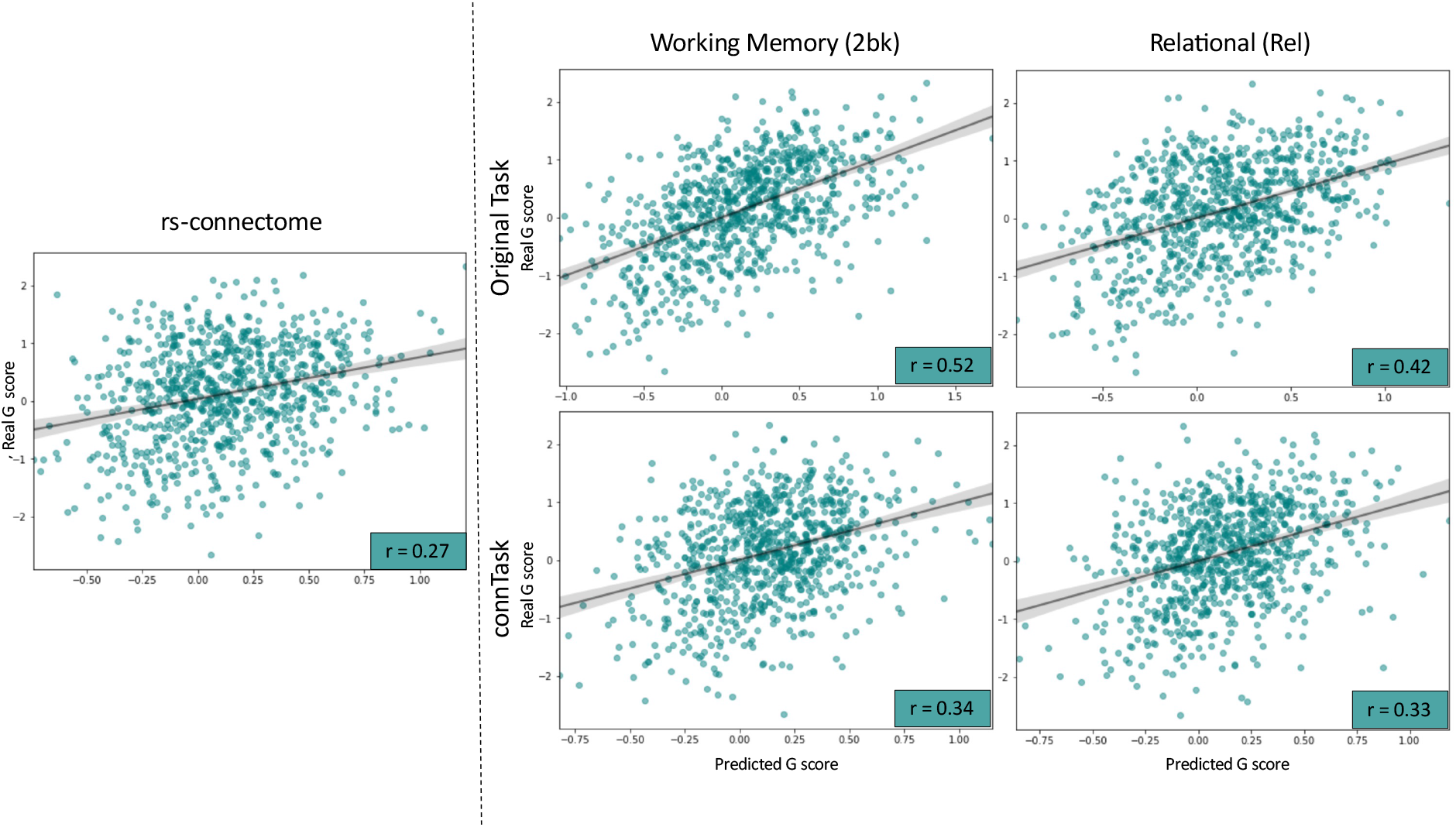
Scatter plots showing G scores prediction success from original and connTask maps of two representative tasks as well as from the rs-connectome. Each point represents a single participant’s real and predicted G scores. Pearson’s r is reported at the bottom-right corner of each plot. Predictions from connTask maps (right, lower panel) were more accurate than those based on the rs-connectome (left panel) but less accurate than predictions based on the original task activation maps (right, top panel)

### Prediction of other individual traits

The same analyses were carried out to predict more specific measures of intelligence, such as reading ability and matrix resolution scores. Additionally, we examined the model’s ability to predict a personality trait, “openness to experience”, the personality dimension best predicted from rs-connectomes (Dubois et al., 2018). Results were similar to the main analysis, showing significantly more accurate predictions by models based on connTask maps rather than on the rs-connectome (V < 0.0001, Bonferroni corrected for 28 comparisons; see Supplementary Figures S5-7).

### Prediction of in-scanner cognitive scores

While the individual traits predicted in this work were collected in tests performed outside of the scanner, it is intriguing to examine our proposed method’s ability to predict scores of tasks performed in-scanner as well.

For this purpose, we used the working-memory task, as this task had a relevant ‘within-scanner’ score. We predicted individual scores from the in-scanner task (N-back task), using models based on the original and connTask maps of this task (specifically, the ‘2bk’ contrast), as well as a model based on the resting-state connectome. In line with the main analysis results, predictions of the N-back task scores were significantly more accurate when using the connTask maps than the rs-connectome *(p* < 0.0001). Interestingly, however, the amount by which the model based on the original activation maps outperformed the model based on connTask maps was significantly larger for the in-scanner measurement than for the out-of-scanner measurements (p < 0.0001, Figure 4). This difference between in- and out-of-scanner measurements was also evident in a significant difference in the ratio of prediction success between original and connTask maps, for G-scores and reading abilities prediction (p < 0.0001; Figure 4).

**Figure 4.**
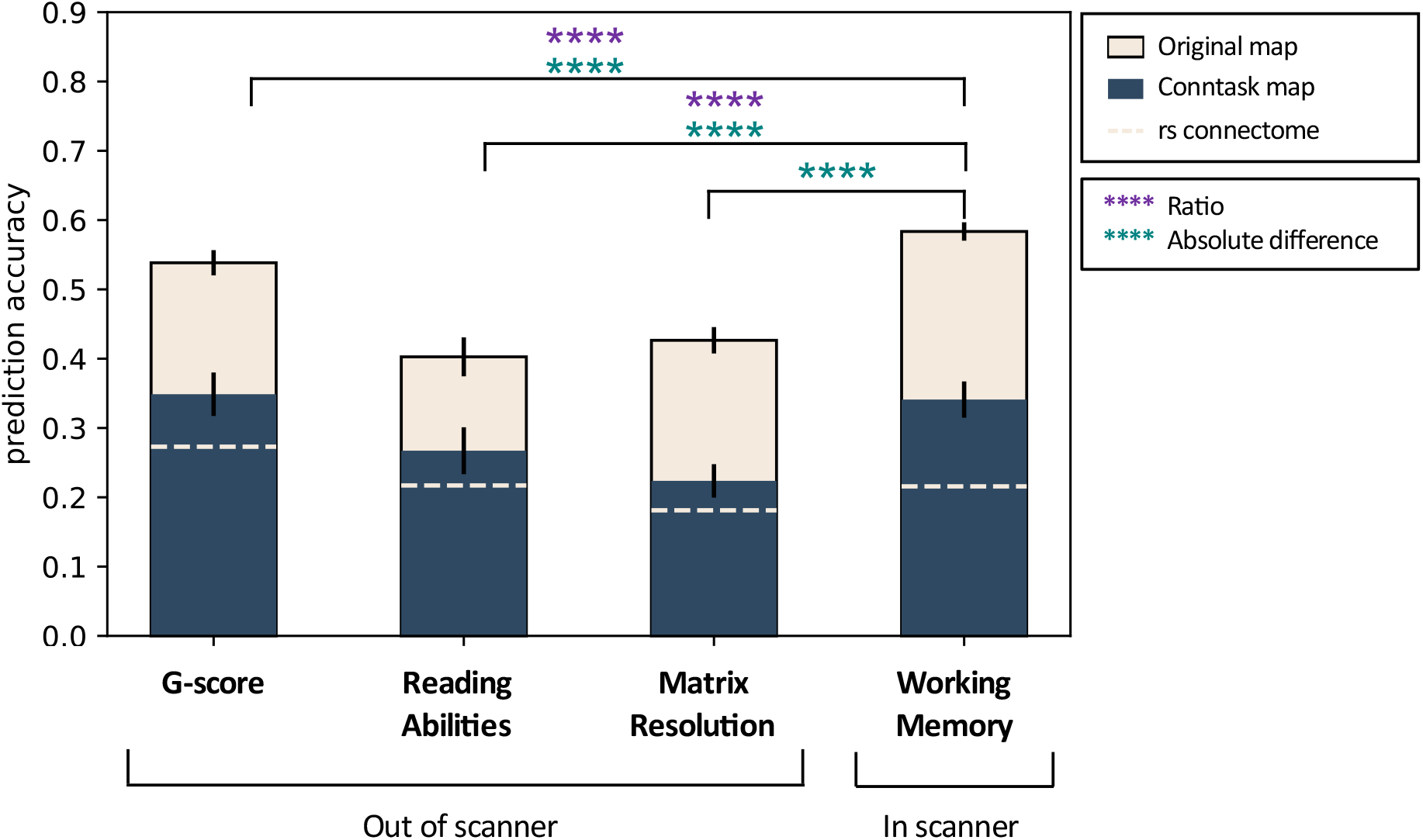
Prediction of in-scanner and out-of-scanner measurements using models based on original and connTask maps of the Working Memory task, and the rs-connectome. Error bars depict standard error of the prediction success between cross-validation folds. Significance asterisks (**** *p* < 0.0001) relate to either the absolute differences (teal) or the differences in ratio of prediction success (purple) between predictions made from original and connTask maps. The difference between connTask and rs-connectome based predictions was significant as well, for all the measurements shown here (p < 0.0001).

### Investigating the predictive power of different brain networks

To explore the contribution of different brain networks to the G-score prediction from each (actual or predicted) task contrast, we generated maps that quantify each vertex’s contribution to the prediction model (see methods and Supplementary Figure S8). We then counted the number of significantly contributing clusters in each resting-state network (Yeo et al., 2011). The largest number of contributing clusters was found in the frontoparietal, attention and default mode networks, indicating that these networks highly contributed to prediction success from both original and connTask activation maps (Supplementary Figure S8). The contribution of these networks to the G-score prediction was independent of task-activation prediction accuracy (Supplementary Figure S12).

## Discussion

Our findings demonstrate a successful prediction of a complex cognitive trait, general intelligence, as well as of more specific cognitive abilities (e.g., reading, matrix resolution) and even personality dimensions (e.g., openness), from brain function at rest. While most methods for predicting individual traits from rs-fMRI are based on transforming the rs-fMRI time-series to a parcellated connectivity matrix (i.e., the rs-connectome), the prediction presented here was achieved by representing the multi-dimensional resting-state data as task-induced brain-activation maps of unperformed tasks. Predictions of individual traits using models based on these maps were significantly better than predictions from models based on the rs-connectome. Our findings thus suggest that the standard connectome may not be the best representation of the resting-state signal for the purpose of predictive modeling, and encourage future studies to explore alternative representations.

Brain areas in the frontoparietal, attention and default mode networks contributed the most to the prediction of general intelligence scores. The contribution of these networks to G-score prediction was independent of the accuracy of task-activation prediction (Figure S12) and was also evident in G-scores prediction from the original activity maps (Figure S8). This finding is in line with previous literature linking the frontoparietal and default mode networks to individual differences in intelligence. Specifically, resting-state functional connectivity within and across the frontoparietal and default mode networks has been positively correlated with G-scores in a cohort of 317 unrelated HCP participants (Hearne et al., 2016). Consistently, differential activation of these networks during cognitive tasks performance was associated with inter-individual differences in intelligence scores (Basten et al., 2013). Our findings therefore add to the accumulating evidence regarding the brain networks that support human intelligence.

In addition to the predictions of general intelligence scores, reading ability, matrix resolution and openness to experience, which are all based on tests performed outside the MRI scanner, we tested our proposed approach’s ability to predict scores obtained *during* an fMRI scan, i.e., the working memory (N-back) scores. While our results confirmed that connTask-based models outperform connectome-based models in predictions of in-scanner measurements, they also revealed a larger gap in prediction success between connTask and original task-based models, compared to this gap in prediction of out-of-scanner measurements.

This finding points to two factors that may influence performance of cognitive tasks: first, participants’ general cognitive ability, which is a rather fixed trait within an individual, and second, the current cognitive state during task performance. Whereas both factors may be represented in data acquired *during* task performance, resting-state data only capture participants’ traits (Gratton et al., 2018). Therefore, predictions of in-scanner tasks may be more accurate when based on data that reflect both the trait and state of the task-performing participants. A trait vs. state representation may also offer an explanation as to why predictions of the global G factor were actually higher when using the connTask maps as opposed to the original task activation maps in a large number of participants (see Supplementary figures S3-4): it is possible that for those participants, the volatile state while performing the task was actually a disturbance for the prediction of a general trait.

We were able to accurately predict activity maps for a diverse selection of tasks and demonstrated that models based on combinations of task-activation maps are more accurate than those based on a single map. In fact, the combination of multiple contrast maps improved not only the prediction of traits from connTask maps but also from the actual activity maps (see Figure 2A), providing a high accuracy of 0.62 for G-scores prediction. While the use of actual activation maps for the purpose of trait prediction is preferable, it is not always feasible, such as in the case of non-compliant populations, or existing datasets that do not include the task of interest. In such cases, prediction from connTask maps may be a promising alternative.

The resting-state signal offers a considerable amount of information from which features for predictive modeling can be extracted by various statistical approaches. Prediction of individual task-activation maps from rs-fMRI, as we propose here, serves as a novel method for feature extraction. Even though there may be a specific, carefully designed fMRI task that could generate a better prediction for each trait than resting-state derived data, the approach we suggest provides the opportunity to produce activation maps for many different tasks at once, even tasks not included in the original dataset. This notion highlights the potential of our method to evaluate measures of human behavior and cognitive abilities, from a variety of (predicted) brain activity patterns, using data from a simple fMRI design.

Moreover, since it does not require participants to actually perform any task, our method enables the study of unique populations, such as psychiatric or neurologic patients, which might not be able to comply with in-scanner tasks. Several studies have already demonstrated successful predictions of task-induced activity patterns in such populations (Mill et al., 2020; Parker Jones et al., 2017; Tik et al., 2021), and in larger datasets, relating these predicted activity maps to behavioral and cognitive traits, or even symptoms, is a promising possibility.

Given that our connTask maps were predicted from resting-state connectivity, one may wonder how could they produce an improved prediction of cognitive traits relative to the original restingstate connectivity patterns. The additional information, beyond connectivity patterns, that enabled this improvement in prediction may lie in the task-prediction model itself, which was trained to relate between brain connectivity and activity. Thus, connTask activity maps serve as a novel representation of resting-state connectivity data, in which the vast data within resting state fMRI is reduced to a form that may be more appropriate for the study of brain-behavior associations (e.g., functional connectivity fingerprinting).

Several limitations of this work that should be discussed concern the data used for our methods’ development and testing, taken from the Human Connectome Project 1200 subjects release. First, this dataset is unique in terms of acquisition (i.e., high spatial and temporal resolution), the amount of data available for each subject (e.g., 4 resting state scans amounting to a full hour) and processing, and is not representative for standard imaging protocols used in basic and clinical research. Hence, it is important for future studies to test our method performance on standard-quality datasets. Second, while a recent work has suggested that brain-wide association studies should be based on very large amounts of data *(n >* 2000; (Marek et al., 2020)), here, we were limited by the number of HCP participants, and further limited our data pool by only including participants that completed all of the necessary scans and tests that are required for our analysis. It would be beneficial in the future to test this method on larger datasets including thousands of participants and more diverse populations, such as neuropsychiatric patients as mentioned above.

## Conclusion

This work holds valuable impact on both basic research and clinical practice, as it can potentially make the scanning process considerably easier for participants to cooperate with, and could substantially simplify the design of a neuroimaging study. Our purposed method demonstrates the potential and versatility of the resting-state signal. This work emphasizes the extent by which the spontaneous activity of our brain can explain variation in cognitive traits and behavior, that can be uncovered by carefully analyzing this spontaneous activity rather than fitting dedicated tasks to each cognitive trait of interest. In that sense, if we are to borrow the treadmill analogy (Greene et al., 2018; Sripada et al., 2020), we provide a “treadmill test for cognition” without actually walking the treadmill.

## Acknowledgments

Data were provided [in part] by the Human Connectome Project, WU-Minn Consortium (Principal Investigators: David Van Essen and Kamil Ugurbil; 1U54MH091657) funded by the 16 NIH Institutes and Centers that support the NIH Blueprint for Neuroscience Research; and by the McDonnell Center for Systems Neuroscience at Washington University.

The authors acknowledge with thanks the support of the Israel Science Foundation (ISF grant no. 1603/18).

