## supplementary for "Predicting Individual Traits from Unperformed Tasks"

### Supplementary Materials

Shachar Gal[1,2], Niv Tik[1,2], Michal Bernstein-Eliav[1,2], Ido Tavor[*,1,2,3]

[1]Sackler Faculty of Medicine, Tel Aviv University, Tel Aviv, Israel

[2]Sagol School of Neuroscience, Tel Aviv University, Tel Aviv, Israel

[3]Strauss Center for Computational Neuroimaging, Tel Aviv University, Tel Aviv, Israel

[*]**Corresponding Author:**

Dr. Ido Tavor; ORCID iD https://orcid.org/0000-0002-9117-4449

Department of Anatomy and Anthropology

Sackler Faculty of Medicine

Tel Aviv University

Ramat Aviv 6997801

Israel

| Task | Contrast | Diagonal mean | Diagonality index |
|---|---|---|---|
| **Working Memory** | 2bk>0bk | 0.537 | 0.128 |
| | **2bk** | 0.734 | 0.141 |
| | 0bk | 0.716 | 0.117 |
| **Language** | **Math-Story** | 0.741 | 0.16 |
| **Social** | **Random** | 0.724 | 0.119 |
| | TOM | 0.753 | 0.127 |
| | TOM-Random | 0.498 | 0.1 |
| **Relational** | **Rel** | 0.761 | 0.121 |
| | Match | 0.745 | 0.116 |
| **Gambling** | Reward | 0.695 | 0.13 |
| | **Punish** | 0.676 | 0.131 |
| **Emotion** | **Faces-Shapes** | 0.546 | 0.114 |

**Table S1. Prediction success of task-activation maps from resting-state connectivity.** Task-induced brain activation for 14 contrasts from 6 tasks available in the HCP dataset were predicted from resting-state connectivity. For each contrast, we correlated each participant's predicted activation map with the real activation map from all the other participants. We then quantified the prediction success for each contrast using two measurements, following the accuracy estimation used in Tavor et al. (2016).

**Diagonal mean**: the mean correlation across participants between real and predicted activation maps from the same subject. A measure of the prediction's accuracy.

**Diagonality index**: the difference between the diagonal mean and the off-diagonal mean. The off-diagonal mean represents the similarity of the participants predicted map to real maps of all the other participants. A measure of the prediction's specificity.

Highlighted contrasts are the ones plotted in Figure 2 and Supplementary Figures S3-S5.

| Data | Original/connTask | Wilcoxon W | p |
|---|---|---|---|
| **Working-Memory** | original | 500213 | 3.9e-165 |
| | predicted | 400592 | 3.7e-130 |
| **Language** | original | 499010 | 1.4e-163 |
| | predicted | 383412 | 1.9e-48 |
| **Social** | original | 478251 | 8.8e-138 |
| | predicted | 390915 | 8.5e-54 |
| **Relational** | original | 584347 | 2.4e-146 |
| | predicted | 420864 | 3.8e-78 |
| **Gambling** | original | 482075 | 2.3e-142 |
| | predicted | 408080 | 3.5e-67 |
| **Emotion** | original | 441677 | 8.6e-98 |
| | predicted | 394127 | 3.4e-56 |
| **Working-Memory + Emotion** | original | 500105 | 5.4e-165 |
| | predicted | 454470 | 5.4e-111 |
| **Working-Memory + Relational** | original | 500488 | 1.7e-165 |
| | predicted | 430239 | 1.03e-86 |
| **Working-Memory + Language** | original | 500497 | 1.6e-165 |
| | predicted | 411287 | 7.5e-70 |
| **Language + Relational** | original | 500435 | 2e-165 |
| | predicted | 433500 | 8.4e-90 |
| **Language + Relational + Emotion** | original | 500423 | 2e-165 |
| | predicted | 458981 | 7.6e-116 |
| **Working-Memory + Language + Relational** | original | 500483 | 1.7e-165 |
| | predicted | 432646 | 5.4e-89 |
| **Working-Memory + Language + Social + Emotion** | original | 500480 | 1.7e-165 |
| | original | 474026 | 8.3e-133 |
| **Working-Memory + Language + Social + Emotion + Gambling +Relational** | original | 500420 | 2e-165 |
| | predicted | 467060 | 8.3e-125 |

**Table S2. Wilcoxon rank test scores for all prediction models.** Detailed results of all comparisons between models based on activation maps (predicted/original) and models based on the resting state connectome. Significance testing procedure is described in the methods section.

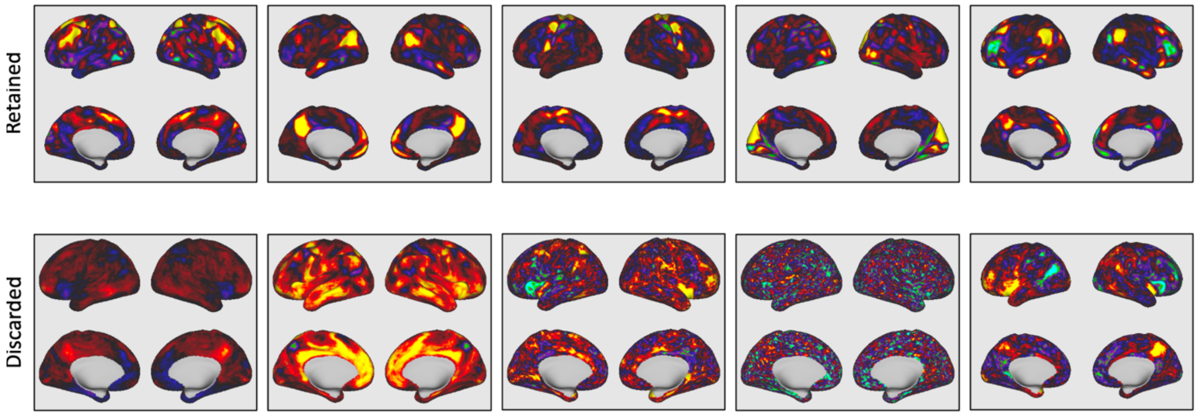


**Figure S1. Examples of independent components yielded in the group-ICA step of the connTask generation pipeline** (see methods). The top row shows components that were included in the subsequent analyses. The bottom row shows the 5 components that were dropped from all subsequent analyses, due to their classification as noise, or their signal originating in areas where there is high CSF contamination.

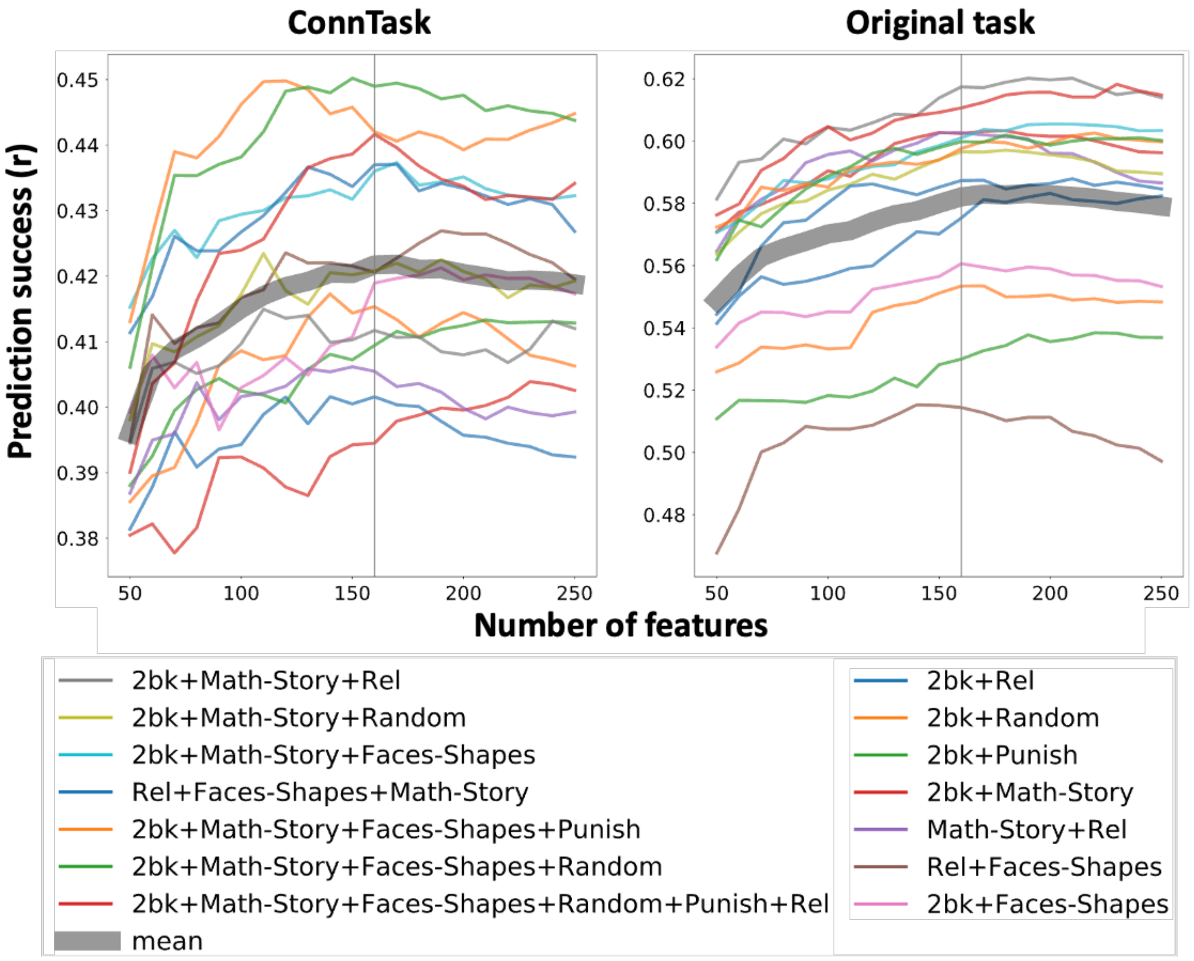


**Figure S2. Examination of a varying number of features in the multi-image BBS pipeline.** We examined the effect of the number of features used in the expanded BBS pipeline on prediction success. Each line plots prediction success for a model based on a specific combination of connTask maps, using a varying number of features. The gray line represents the mean prediction success of all models across all numbers of features, and is plotted to illustrate the general trend. For the main analysis we chose to use 160 features, which is the optimal number for the connTask combinations, and close to optimal in the original task maps.

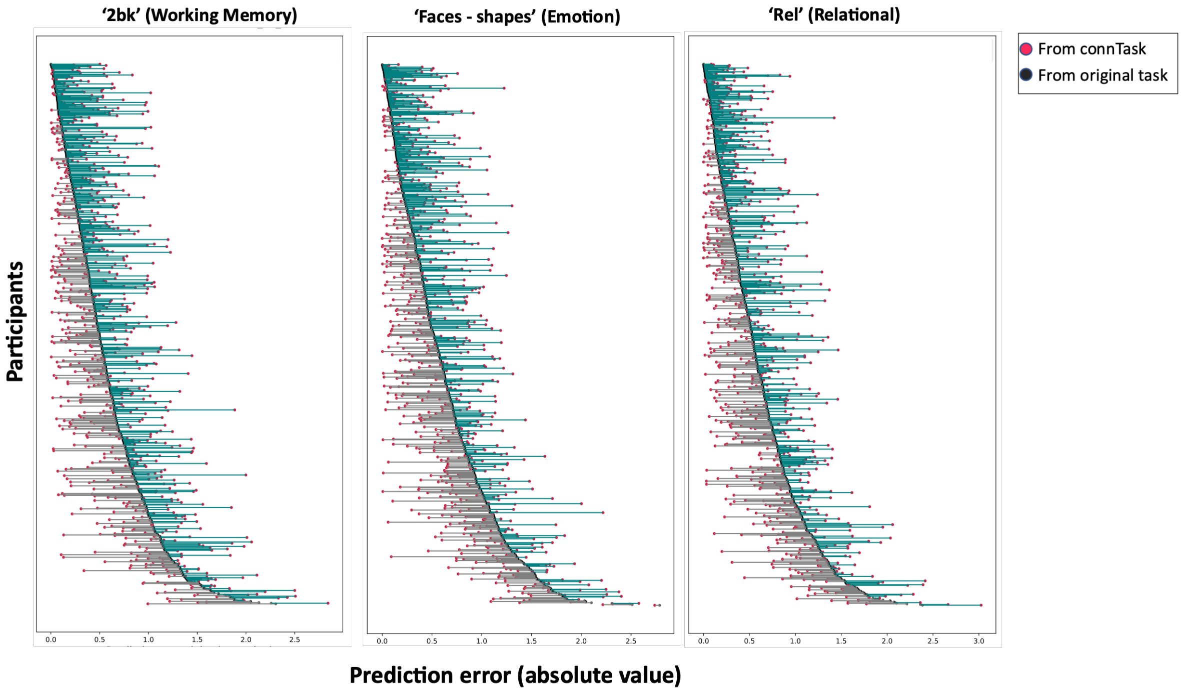


**Figure S3. Comparison of individual prediction errors between original and predicted task-activation maps.** Even though prediction success was better, on the group average, for models based on original task-activation maps than those based on connTaks maps, we sought out to examine this effect on an individual basis. For each participant, in each contrast, we calculated the absolute error between actual and predicted G-scores, based on either the original task-activation map or the connTask map. The figure presents these individual absolute errors in three exemplary task contrasts.

The data is sorted by the absolute error of predictions yielded from original task activations (black dots). The red dots represent the absolute error of predictions based on connTask maps. A horizontal line is plotted to connect each participants' errors, colored in gray or teal to emphasize the fact that there was a large portion of participants where the prediction error of the connTask-based model was smaller than the prediction error of an original task-based model. As shown in the figure, the red dots appear either on the left side or on the right side of the black ones, indicating individual variability: for some of the participants, prediction was better from connTask than from original task maps (red dot left to the black one, connected by a gray line), while for other participants, prediction was better from the original than from connTask maps (red dot right to the black one, connected by a teal line).

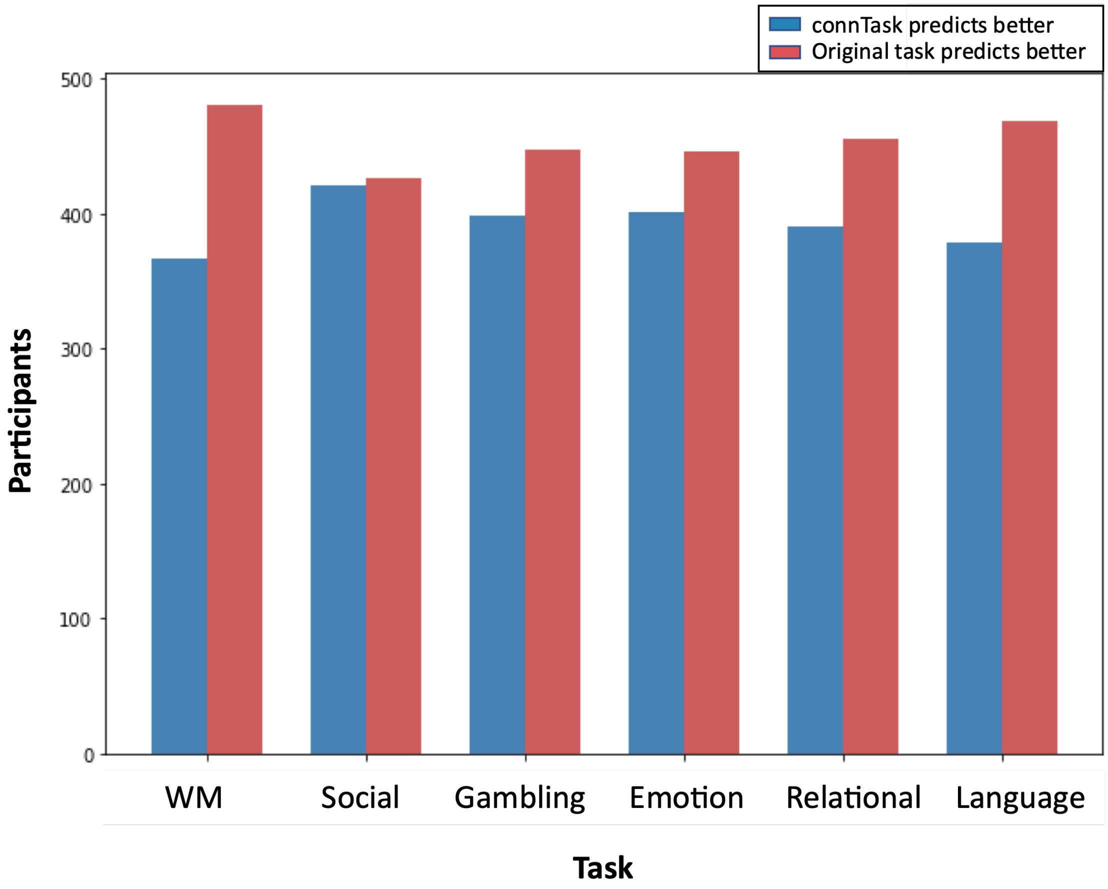


**Figure S4. Distribution of participants showing better prediction of G-score from original (in red) or predicted (in blue) task-activation maps.** In all task contrasts the majority of participants' G-scores were better predicted using original task maps than connTask maps. However, for each contrast there is a large subset of participants for which predictions based on the connTask maps were more accurate, as shown in the figure for an exemplary contrast for each task.

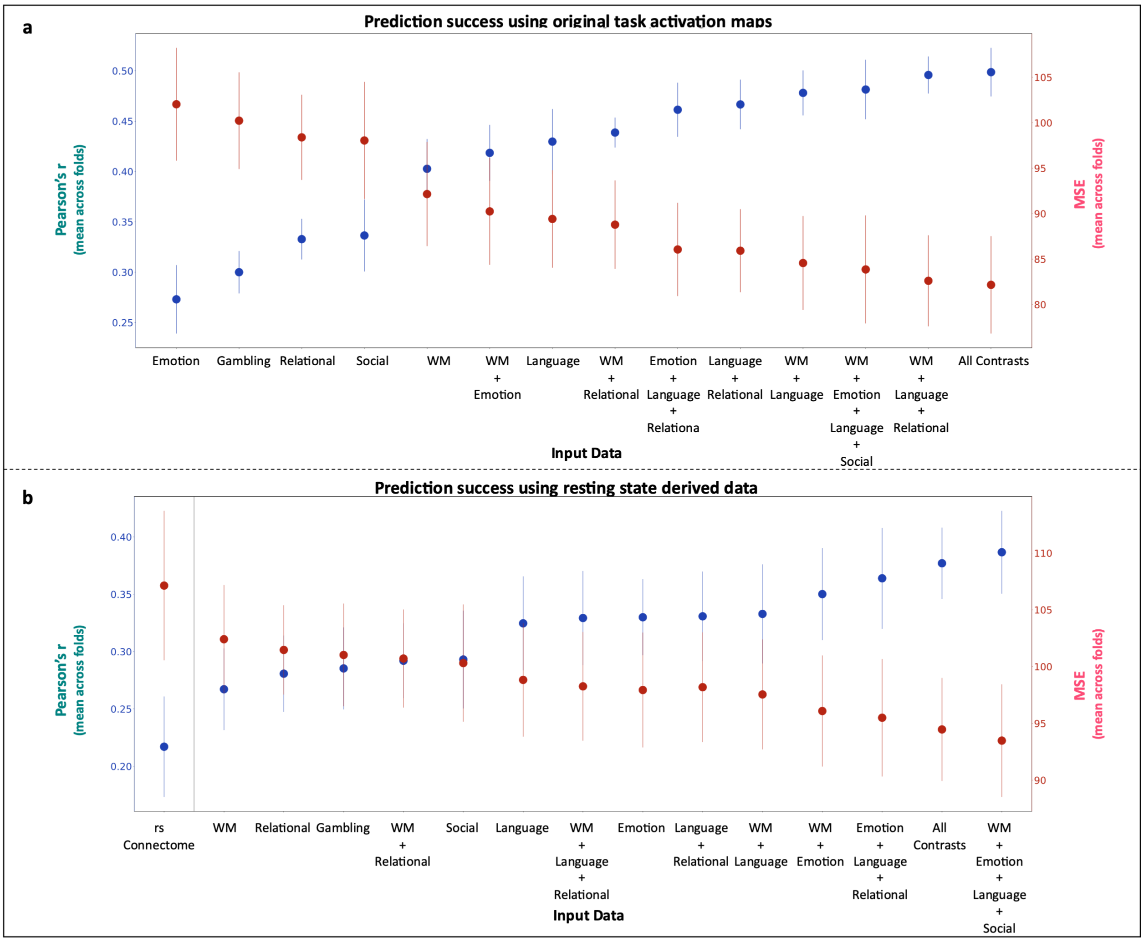


**Figure S5. Reading abilities prediction results**. The figure presents prediction results of the ReadEng_Unadj score, which reflects reading abilities and is included in the G-score calculation. Prediction success was estimated by the Pearson correlation between predicted and observed scores (in green, referring to the left y-axis) and the mean squared error of the prediction (in red, referring to the right y-axis). A) Predictions from real task activation maps. B) Predictions from resting-state derived data (connTask maps and resting-state connectome). All the predictions based on connTask maps are were significantly more accurate than the prediction based on rs-connectome ($p < 0.0001$).

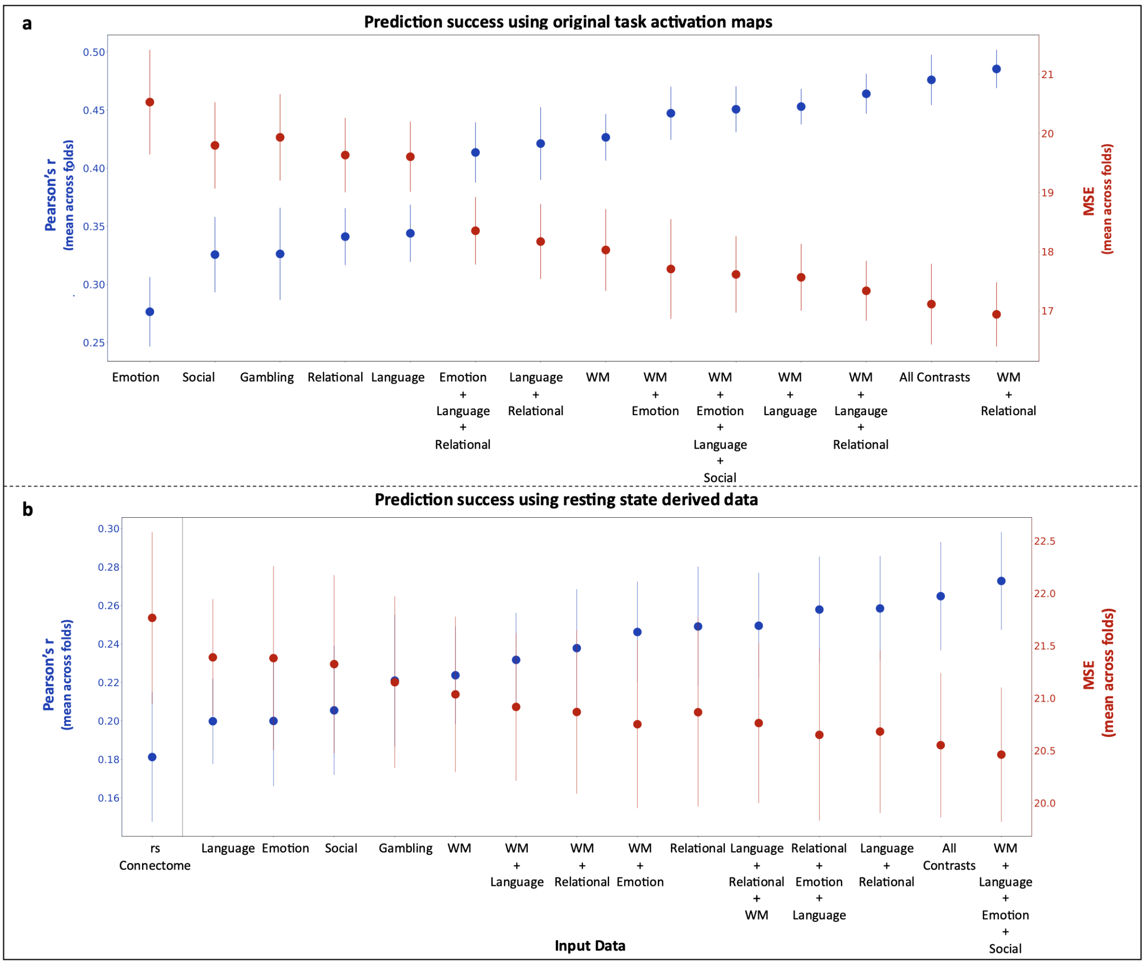


**Figure S6. Matrix resolution prediction results**. The figure presents prediction results of the Pmat_24_CR score, which reflects matrix resolution abilities, and is included in the G-score calculation. Prediction success was estimated by the Pearson correlation between predicted and observed scores (in green, referring to the left y-axis) and the mean squared error of the prediction (in red, referring to the right y-axis). A) Predictions from real task activation maps. B) Predictions from resting-state derived data (connTask maps and resting-state connectome). All the predictions based on connTask maps are were significantly more accurate than the prediction based on rs-connectome ($p < 0.0001$).

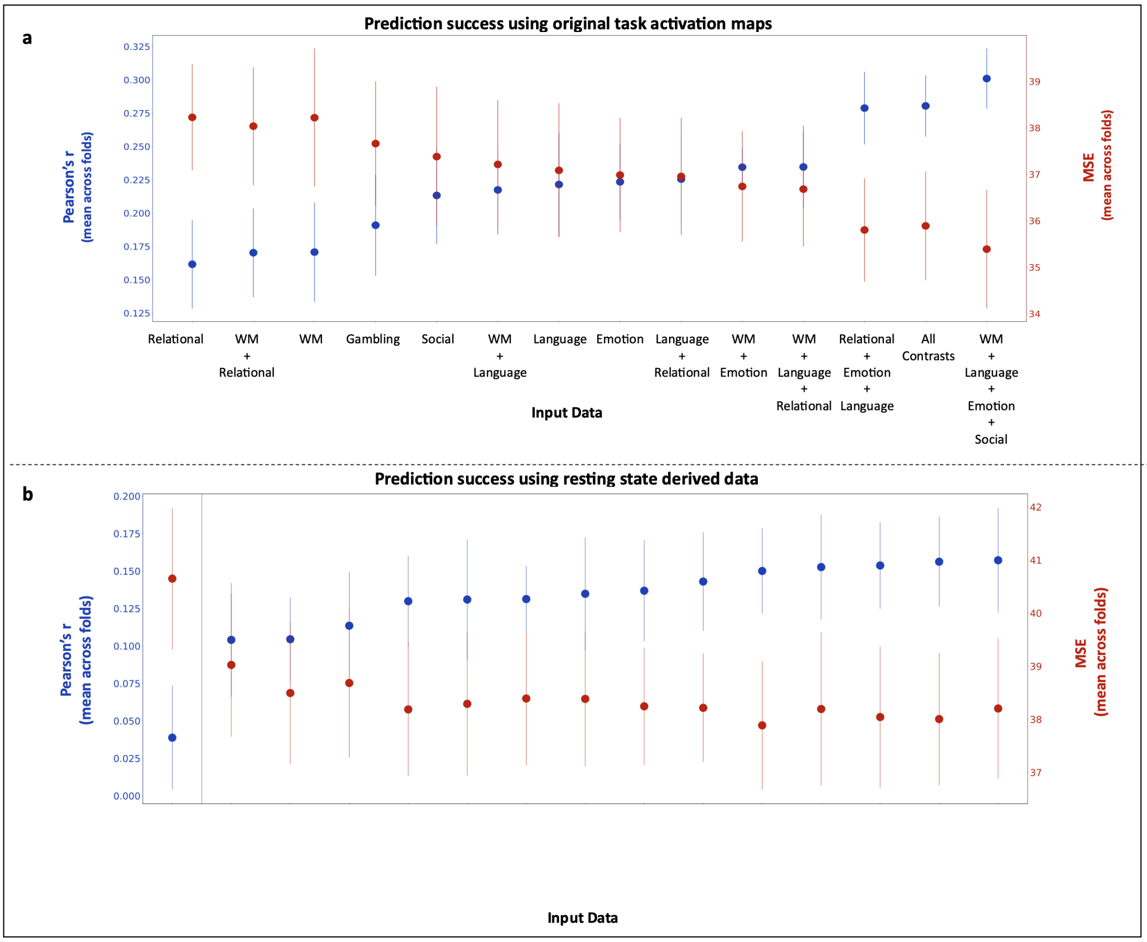


**Figure S7. "Openness to experience" prediction results.** The figure presents prediction results of "Openness to experience". Of the five personality dimensions described in the "Big-5" theory (McCrae and Costa, 1987; McCrae and John, 1992) "Openness to experience" is the dimension best predicted from rs-connectomes (Dubois et al., 2018). Prediction success was estimated by the Pearson correlation between predicted and observed scores (in green, referring to the left y-axis) and the mean squared error of the prediction (in red, referring to the right y-axis). A) Predictions from real task activation maps. B) Predictions from resting-state derived data (connTask maps and resting-state connectome). All the predictions based on connTask maps are were significantly more accurate than the prediction based on rs-connectome ($p < 0.0001$).

In a post-hoc analysis, we tested the prediction of the remaining 4 personality dimensions (conscientiousness, extraversion, agreeableness and neuroticism). Results from all input types (original/predicted task activation maps and rs-connectomes) were generally low, and other than "Extraversion", none showed significantly improved predictions from connTask data, compared with rs-connectome.

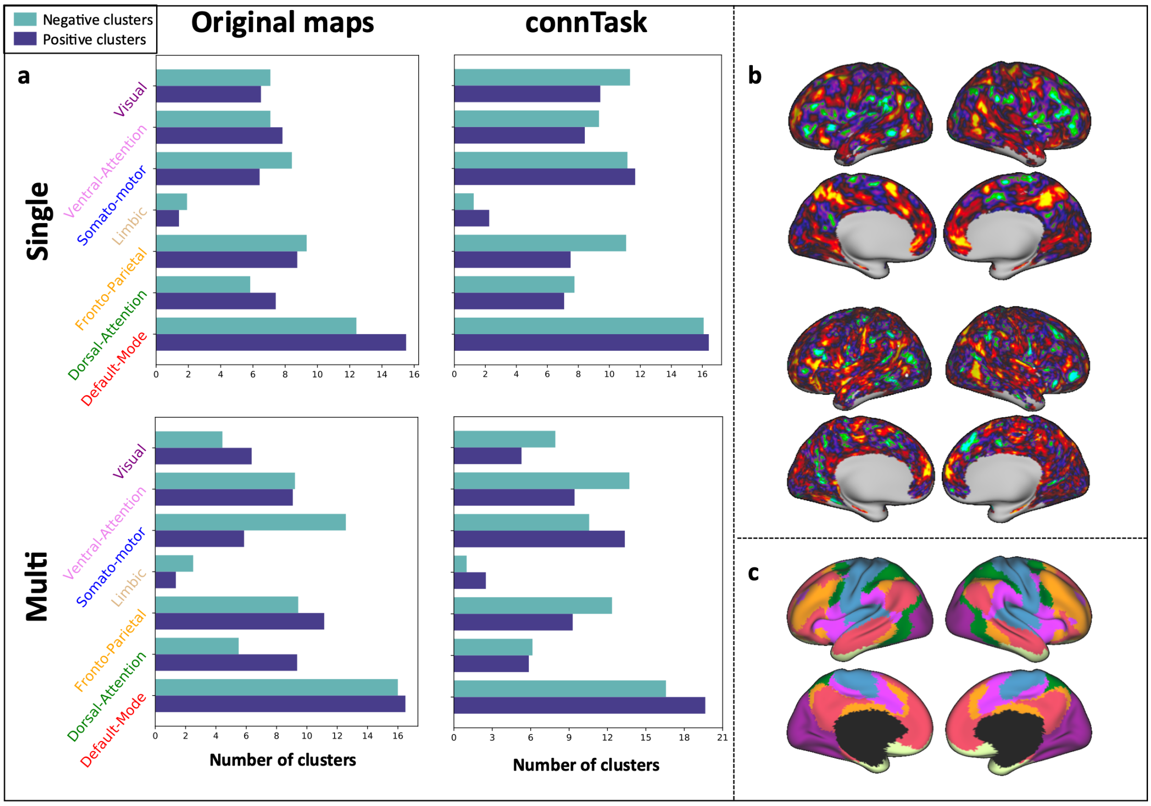


**Figure S8. Clusters of significant contribution to prediction, divided by resting-state networks**. We created vertex-wise maps of contribution to prediction and calculated the number of predictive clusters (cluster size > 15, vertex contribution value > 0.99 percentile) in each resting-state network (Yeo et al., 2011). (**a**) Number of clusters of significant contribution to prediction within each resting-state network. Top row represents the contribution to prediction from a single contrast map, either original (left) or predicted (right), averaged across task contrasts. Bottom row represents the contribution to prediction from multiple original (left) or predicted (right) contrast maps, averaged across contrast combinations. (**b**) Examples of maps depicting vertex-wise contribution to the prediction, using the original (top) and connTask (bottom) maps of the '2bk' (working memory) contrast. (**c**) The division of the cortex to 7 resting-state networks suggested by Yeo et al. (2011), which was used for the analysis in panel **a**.

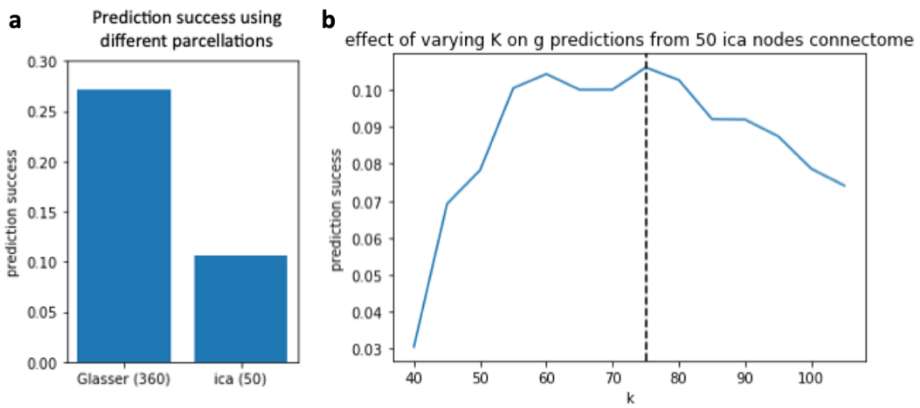


**Figure S9. Prediction success using different parcellations.** In our main analysis, a 360-node parcellation (Glasser et al., 2016) was used to create functional connectomes, whereas a more gross division of 50 nodes, derived from ICA on resting-state data, was used to fit models in the connTask generation pipeline. To avoid biases resulting from the choice of parcellation, we repeated our connectome analysis using the 50-node parcellation and predicted g-scores using the single-map BBS pipeline. **(a)** Prediction accuracy (calculated as the Pearson's correlation between real and predicted scores) was 0.106, which is considerably lower than prediction accuracy from connectomes built on the 360 nodes parcellation. This result suggests that 50 nodes, resulting in 1225 edges, is probably not a rich enough representation of connectivity data to be used in a BBS paradigm. **(b)** We tested different values for the parameter K in the BBS paradigm. We found that K=75 is still the optimal value even with the suboptimal connectomes based on the 50-node parcellation.

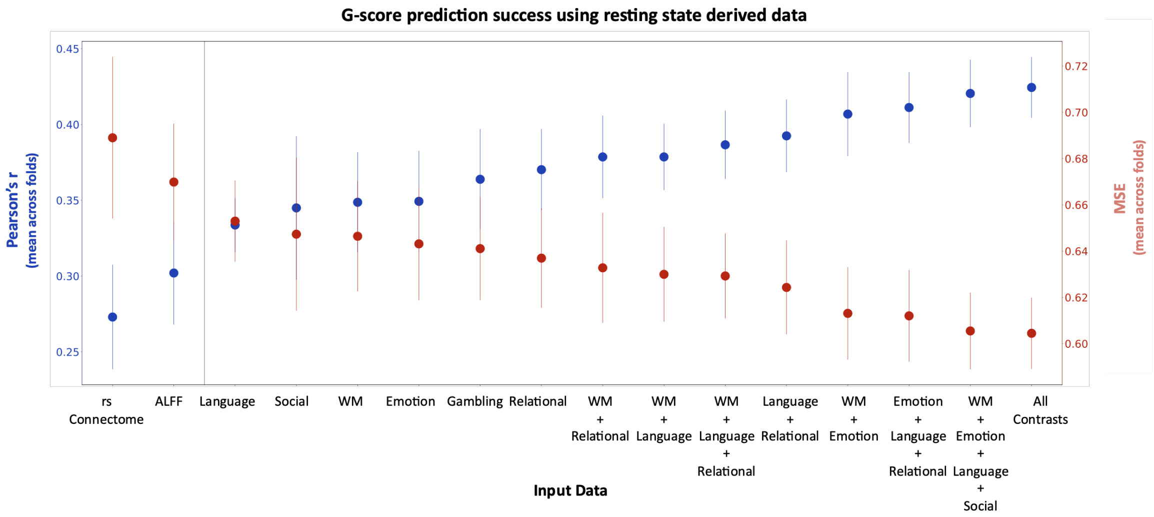


**Figure S10. Predictions using connTask maps outperform predictions using ALFF.** In our main analysis, we compared predictions derived from connTask maps with those based directly on the resting-state functional connectome**.** An inherent difference between connectomes and connTask maps is that connectomes depict edgewise data (connectivity) whereas connTask maps depict vertex-wise data (predicted activation). In order to test our method against a more similar, yet less widely used, representation of the resting-state signal, we created maps depicting vertex-wise amplitude of low frequency fluctuations (ALFF), and used them to predict g-scores using the single-map BBS pipeline. While prediction using ALFF maps outperformed the prediction from the resting-state connectome ($p<0.0001$), predictions based on connTask maps were significantly more accurate than those based on ALFF ($p<0.0001$).

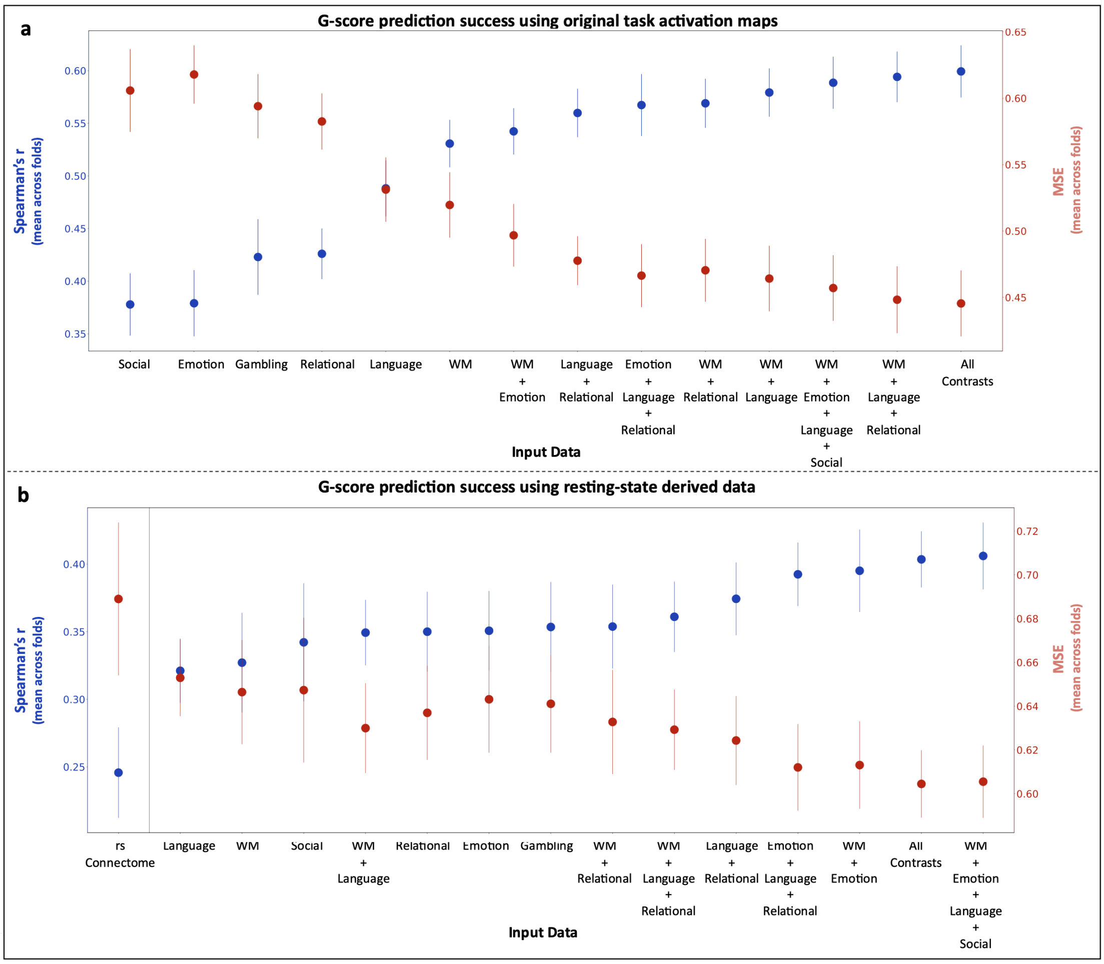


**Figure S11**. **G-score prediction success of models based on real activation maps, predicted activation maps and the rs-connectome.** This analysis is similar to that shown in Figure 2 in the main text, with prediction success calculated as the Spearman's correlation coefficient between real and predicted scores, rather than the Pearson's correlation. The pattern of results remains similar, showing more accurate predictions based on original data than on connTask data **(a)** and more accurate predictions derived from connTask data than from connectome data **(b)**.

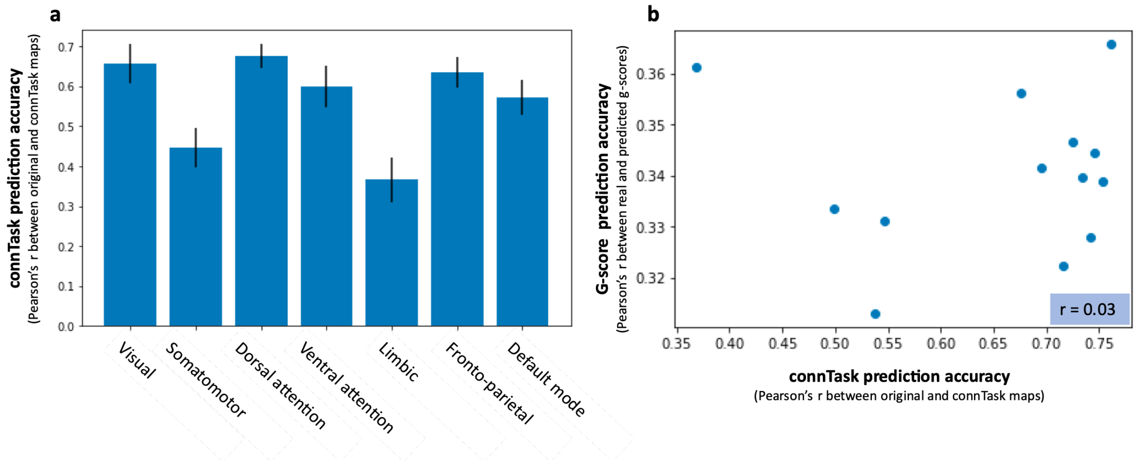


**Figure S12. Accuracy of trait prediction is not dependent on connTask accuracy. A)** The accuracy of the prediction of task-activation patterns (averaged across tasks) in each resting-state network separately. A comparison with Figure S8 suggests that there is no dependency between the accuracy of activation prediction in each of the networks and the contribution of these networks to the g-prediction models (e.g., prediction of task-activation was high in the visual network, but its contribution to the prediction of intelligence was relatively low). **B)** A scatter plot showing no correlation between the accuracy of task-activation prediction (each task is represented with a blue dot) and the accuracy of G-scores prediction from this task contrast, suggesting that the success of G-score prediction from connTask maps is not dependent on connTask maps accuracy.
